# Aging amplifies a gut microbiota immunogenic signature linked to heightened inflammation

**DOI:** 10.1101/2024.03.22.586320

**Authors:** Maria Elisa Caetano-Silva, Akriti Shrestha, Audrey F. Duff, Danica Kontic, Patricia C. Brewster, Mikaela C. Kasperek, Chia-Hao Lin, Derek A. Wainwright, Diego Hernandez-Saavedra, Jeffrey A. Woods, Michael T. Bailey, Thomas W. Buford, Jacob M. Allen

## Abstract

Aging is associated with low-grade inflammation that increases the risk of infection and disease, yet the underlying mechanisms remain unclear. Gut microbiota composition shifts with age, harboring microbes with varied immunogenic capacities. We hypothesized the gut microbiota acts as an active driver of low-grade inflammation during aging. Microbiome patterns in aged mice strongly associated with signs of bacterial-induced barrier disruption and immune infiltration, including marked increased levels of circulating lipopolysaccharide (LPS)-binding protein (LBP) and colonic calprotectin. *Ex vivo* immunogenicity assays revealed that both colonic contents and mucosa of aged mice harbored increased capacity to activate toll-like receptor 4 (TLR4) whereas TLR5 signaling was unchanged. We found patterns of elevated innate inflammatory signaling (colonic *Il6, Tnf, Tlr4*) and endotoxemia (circulating LBP) in young germ-free mice after 4 weeks of colonization with intestinal contents from aged mice compared with young counterparts, thus providing a direct link between aging-induced shifts in microbiota immunogenicity and host inflammation. Additionally, we discovered that the gut microbiota of aged mice exhibited unique responses to a broad-spectrum antibiotic challenge (Abx), with sustained elevation in *Escherichia* (Proteobacteria) and altered TLR5 immunogenicity 7 days post-Abx cessation. Together, these data indicate that old age results in a gut microbiota that differentially acts on TLR signaling pathways of the innate immune system. We found that these age-associated microbiota immunogenic signatures are less resilient to challenge and strongly linked to host inflammatory status. Gut microbiota immunogenic signatures should be thus considered as critical factors in mediating chronic inflammatory diseases disproportionally impacting older populations.

## INTRODUCTION

The gut microbiota changes throughout the lifespan, and especially during older age. Emerging evidence indicates that changes in the microbiota may influence the aging process (DeJong et al., 2020), and is associated with the development and progression of numerous chronic, age-related inflammatory diseases including type 2 diabetes, cancer, and neurological conditions including Parkinson’s disease (PD) and Alzheimer’s disease (AD) (Alsegiani & Shah, 2022; Conway & A Duggal, 2021; Ghosh et al., 2022; Zhu et al., 2021). Conversely, gut microbial signatures may be predictors of healthy aging, including evidence showing that gut microbiota composition and associated metabolic signatures can be used as strong predictors of longevity in humans (Wilmanski et al., 2021). However, how the gut microbiota directly contributes to long-term health and immune system function with advanced age is not fully understood.

Chronic inflammation is a hallmark of both aging and aging-related diseases. The gut microbiota is tightly linked to inflammatory tone due to its plethora of immunogenic signals, including pathogen-associated molecular patterns (PAMPS), including lipopolysaccharide (LPS) and flagellin, that directly activate highly conserved innate Toll-like receptor (TLR) pathways in the host, TLR4 and TLR5, respectively. Under certain environmental challenges, the gut microbiota become dysbiotic whereby gut microbiota PAMPs can directly contribute to unabated inflammation, within the gut and systemically (Graham & Xavier, 2023; Malik et al., 2023). This effect has been proposed as a mechanism in aging populations who exhibit altered gut microbiome composition together with elevated levels of inflammatory cytokines (Thevaranjan et al., 2017). However, questions remain about how aging alters gut microbiota immunogenic patterns that directly facilitate low-grade inflammation.

Older adults frequently experience unique environmental challenges that may further exacerbate shifts in gut microbiome function, potentially increasing the risk for chronic disease. These challenges include antibiotic exposure which is highly prevalent among older adults. For instance, in 2014, it was estimated that U.S adults > 65 years of age received more than 50 million antibiotic courses (Kabbani et al., 2018). Alongside off-target effects and long-term community risk of antibiotic resistance, overuse of antibiotics in older adults may drive long-term detrimental changes to the gut microbiota. This is supported by data indicating that the microbiota of aged mice fail to return to baseline composition for six (6) months following a single course of broad-spectrum antibiotic treatment (Laubitz et al., 2021). However, the functional changes to the microbiome caused by late-life antibiotic use are largely underexplored.

In this study, we determined whether microbiota signatures associated with advanced age harbor a greater capacity to activate host inflammatory signaling using *ex vivo* TLR reporter cell lines and microbiota transplants into previously germ-free animals. We found that colonic contents and mucosa of aged mice exhibit heightened TLR4-signaling that paralleled multiple signs of low-grade intestinal inflammation. The transfer of microbiota from aged to young germ-free mice partially recapitulated increased colonic inflammation and signs of barrier disruption, thus providing direct evidence for the microbiota as a mediator of intestinal immunity during aging. Lastly, we also determined how aging primes gut microbiota responses to a clinically relevant microbiota challenge—a seven-day course of broad-spectrum antibiotics. We demonstrate that the microbiota from aged mice exhibited altered gut microbiota composition and immunogenic signatures in response to an antibiotic challenge.

## RESULTS

### Gut microbiota composition changes with host age and parallel colonic inflammatory status

We compared the gut microbiome composition of distal colon contents from young (3-4 months) vs aged (19-20 months) C57 Bl/6 mice using 16S rRNA gene sequencing. Aged mice demonstrated a marked shift in microbiome composition when compared to young mice, as evidenced by a significant difference in β-diversity (*p*=0.002) and lower α-diversity (Chao1 *p*=0.009) (**Fig. 1A-B**). Moreover, aging induced alterations in gut microbiota composition at multiple taxonomic levels; for example, we found that *Clostridium*, *Erysipelatoclostridium*, *Mucispirillum*, and *Eubacterium ventriosum* were increased in aged mice, while members of *Anaerostipes*, *Bacteroides*, *Lachnospiraceae*, and *Ruminococcaceae* were diminished in aged mice compared to younger counterparts. Altogether, our data shows that aging is associated with gut microbial alterations in the distal colon (**Fig. 1C**).

**Figure 1.**
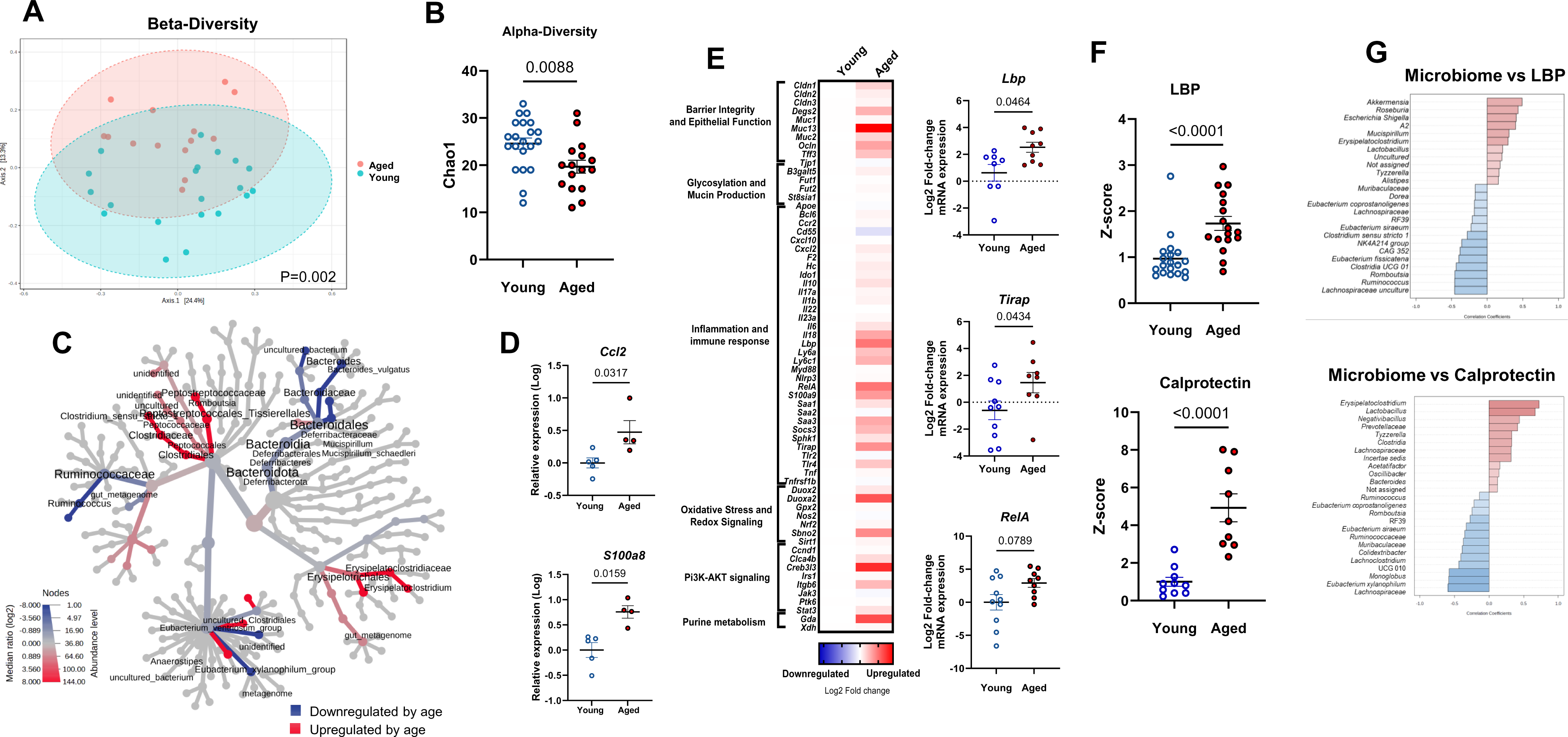
Gut microbiota composition and inflammatory status of young and aged mice. **(A)** β-diversity analysis: Principal Coordinate Analysis plot using the Bray-Curtis index from young (blue spheres) and aged (red spheres) mice. Dotted-line ellipses correspond to clusters in each group. **(B)** α-diversity analysis: index Chao 1 represents community richness. **(C)** Heat tree plotting of colonic taxa abundance (aged vs young) to allow quantitative visualization of community diversity data comparing the two age groups. Microbiome data (Panels A, B, and C) correspond to n= 5 cohorts with n= 15-19 per group. **(D)** Distal colon gene expression in young and aged mice, analyzed via qPCR (n= 4-6 per group). **(E)** Distal colon gene expression in young and aged mice analyzed via Fluidigm and depicted with a heat map. Blue indicates downregulation and red indicates upregulation compared to young mice. Each square represents the mean gene expression per group. Predominantly affected genes by age are represented in individual graphs on the right side of the heat map. Data correspond to n= 5 cohorts with n= 8-15 per group. **(F)** Lipopolysaccharide-binding protein (LBP) in serum and calprotectin in proximal colon contents, both analyzed by ELISA kits according to manufacturer’s recommendations. Values were expressed as Z-score considering young mice as control. Data correspond to n= 3 cohorts with n= 9-18 per group. **(G)** Top 25 genera correlated with LBP and calprotectin expression (Spearman rank correlation). Data correspond to n= 3 cohorts with n= 8-17 per group.

We next determined whether age-dependent alterations in gut microbial composition paralleled host inflammatory status in the distal gut. We identified signs of age-induced innate immune cell infiltration in the colon, including genes encoding the macrophage chemoattractant chemokine ligand 2 (*Ccl2/Mcp1*) and the innate immune cell cytoplasmic protein calprotectin (*S100a8*) (**Fig. 1D**; *p*<0.05). Next, we implemented a custom multiplex Fluidigm gene expression panel that revealed differences in colonic transcription in aged as compared to young mice suggesting innate inflammatory signaling and signs of bacterial penetration into mucosal tissue (**Fig. 1E**). Of note, we found indicators that advanced aged was associated with increased LPS sensing through TLR4, including elevations in colonic expression of LPS-binding protein (*Lbp*) and TIR domain-containing adaptor protein (*Tirap*) (**Fig. 1E**; *p*<0.05). The later encodes a membrane localized adaptor protein that initiates the TLR4 intracellular signaling. Further implicating a heightened innate inflammatory signaling through TLRs, we found a trend for increased Nuclear factor Kappa-B p65 subunit *RelA* in the colon of aged mice (**Fig. 1E**; *p*<0.10). Following up on differences in colonic transcription indicative of bacterial-directed inflammatory signaling, we next determined protein levels of surrogate markers for barrier disruption and innate immune infiltration. We first focused on LBP, given its critical role in the recognition of LPS and modulating the downstream inflammatory cascade. In parallel to expression profiles in the colon, we found significantly higher LBP protein levels in the serum of aged versus their young mouse counterparts (**Fig. 1F**; p<0.001). Next, we measured calprotectin, a critical cytoplasmic protein found in innate immune cells, including neutrophils and macrophages, and a well-known biomarker of immune cell infiltration and intestinal inflammation in inflammatory bowel disease (IBD) patients (Park, 2020; Sakurai & Saruta, 2022). Indeed, we found remarkably higher levels (>4 fold; *p*<0.0001) of calprotectin in colonic contents of aged as compared to young mice. Our data indicates that advanced aged is associated with an increase in innate inflammatory infiltrate within the colonic niche.

To begin delineating potential relationships between age-induced changes in microbiota composition and colonic inflammation, we utilized Spearman rank correlation analysis to reveal multiple age-sensitive bacteria associated with host inflammatory outcomes. For example, *Erysipelatoclostridium* that is increased with age was positively associated with calprotectin and LBP protein expression, whereas members of *Lachnospiraceae* that are decreased by age were negatively associated with both inflammatory markers **(Fig. 1G**; FDR *p*<0.05).

### Microbiota from aged mice possess an increased capacity to activate TLR-4 across multiple colonic regions

Our results indicate that age-induced alterations in colonic microbes correspond to higher colonic and systemic inflammatory markers likely mediated through TLR-signaling, thus we tested the ability of gut microbiota to directly stimulate two canonical innate inflammatory receptors that are sensitive to bacterial PAMPs—TLR4 and TLR5. Using *ex vivo* immunogenicity assays (with reporter cell lines), we found that fecal slurries from aged mice have an elevated capacity to activate TLR4, the canonical receptor for bacterial-derived LPS, compared to their younger counterparts (**Fig. 2B**; *p*<0.0001). To confirm that the phenomenon was not solely associated with the distal region of the gut, we also examined contents in proximal regions of the colon and confirmed that TLR4 reactivity was increased by age in both contents and mucosal scrapings (**Fig. 2C**; *p*<0.05). Unlike TLR4, gut microbiota immunogenicity towards TLR5, the canonical receptor for bacterial-derived flagellin, was unaltered by age in the feces (**Fig. 2D**; *p*>0.05) and proximal colonic regions (**Fig. 2E**; *p*>0.05). This indicates that age-induced alterations in colonic microbes can directly drive inflammation through a mechanism involving TLR4.

**Figure 2.**
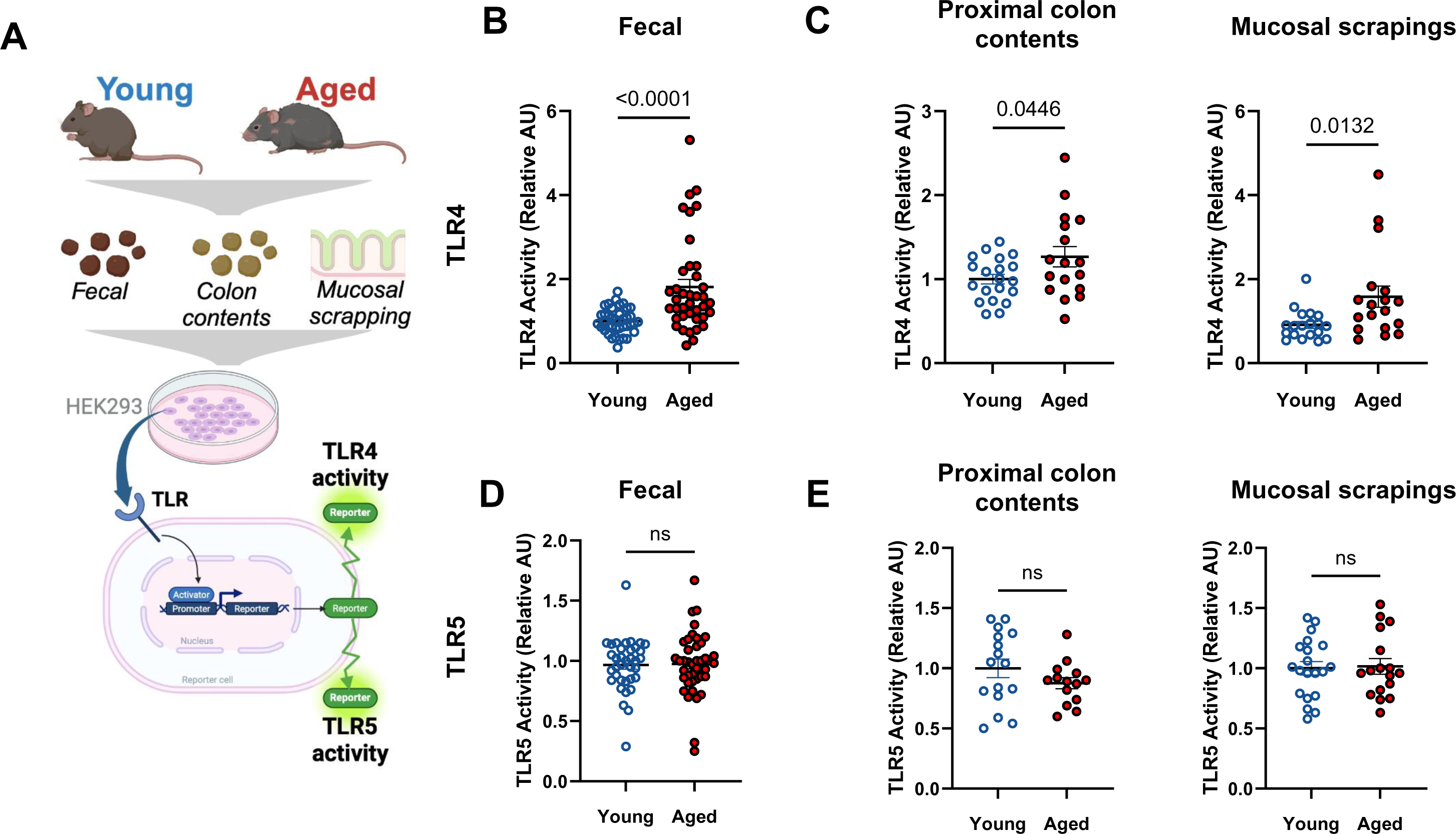
Colonic microbiota from aged mice exhibits elevated capacity to activate TLR4. **(A)** A schematic representation of immunogenicity experiment. The ability of colonic microbiota to activate **(B-C)** TLR4 and **(D-E)** TLR5 activity in **(B and D)** feces and **(C and E)** proximal colon contents and mucosal scrapings, was assessed using human embryonic kidney (HEK293)-Blue-mTLR4 and HEK-Blue-mTLR5 cells. Relative fluorescence intensity using a nuclear-factor Kappa Beta-linked inducible reporter (secreted embryonic alkaline phosphatase) was used to assess TLR4 or TLR5 activity. Data was normalized within each independent experiment and relativized to young control at AU =1. Fecal data correspond to n= 4 cohorts with n= 23-25 per group. Proximal colon content and mucosal scrapings data correspond to n= 2 cohorts with n= 17-21 per group.

### Microbiota transplant from aged mice to young germ-free mice partially recapitulates colon inflammation and signs of intestinal barrier disruption

Following up on *ex vivo* immunogenicity assays, we next wanted to further explore the hypothesis that an age-induced increase in microbiota TLR4 immunogenicity could directly mediate low-grade inflammation *in vivo*. To accomplish this, a mixture of intestinal contents (cecal and colonic contents, and feces) from aged or young donor mice was weighed, homogenized in a single slurry, and transplanted into young gnotobiotic recipients. After 4 weeks of colonization, we found increased markers of colonic inflammation and low-grade endotoxemia in mice that received a microbiota transplantation (MT) from aged donors (**Fig. 3**). In the colon, we found increased expression of the inflammatory genes *Il6, Tnf*, and *Tlr4* in mice that received an aged microbiota transplant (**Fig. 3B**; *p*<0.05) and a trend for increased circulating LBP (**Fig. 3C**; *p*<0.1). Despite confirming that microbiota TLR4 reactivity was increased in aged donor intestinal contents, we found no differences in the TLR4 reactivity in feces (**Fig. 3D**; *p*>0.05) among recipient gnotobiotic animals at 4 weeks post-transfer. These findings suggest age-dependent increases in microbiota TLR4 immunogenicity do not persist after extended periods of colonization in recipients and are likely sensitive to the age of the host.

**Figure 3.**
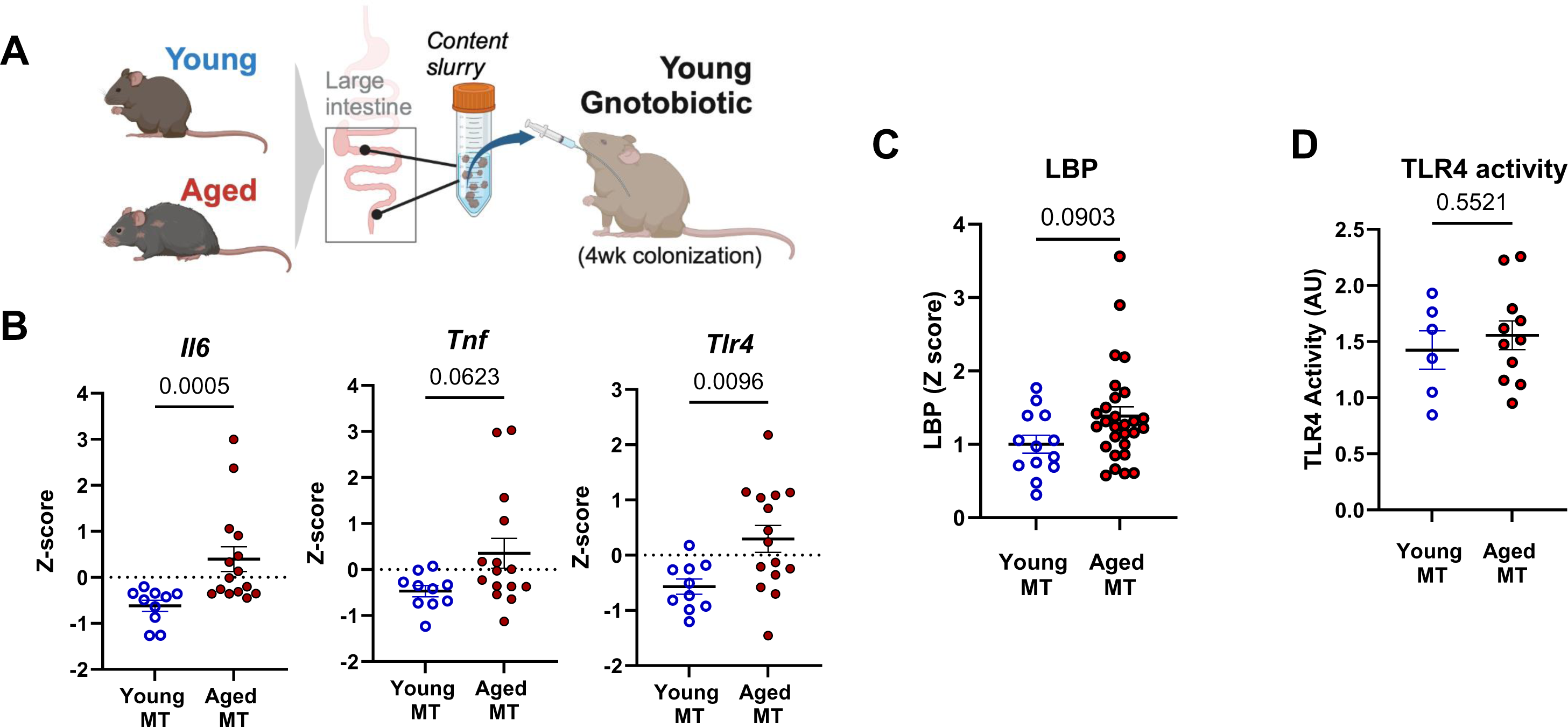
Aged microbiota transplantation (MT) into germ-free mice partially recapitulates colon inflammation and signs of barrier disruption. **(A)** A schematic representation of the gnotobiotic experiment. A mixture of intestinal contents (cecal and colonic contents, and feces) from young (3-4 months) or aged (18-20 months) CONV-R donor mice was transferred to young germ-free recipient mice. All measurements were performed in young MT and aged MT samples, 4 weeks after colonization. **(B)** Colonic inflammatory gene expression of recipient gnotobiotic mice was assessed by qPCR. **(C)** Lipopolysaccharide-binding protein (LBP) in serum, analyzed by ELISA kit according to manufacturer’s recommendations. **(D)** TLR4 activity in colon contents, assessed using human embryonic kidney (HEK)-Blue-mTLR4. Relative fluorescence intensity was used as an estimation for TLR4 activity, considering gnotobiotic mice plus young MT as control (Relative AU =1). Data correspond to n= 3 cohorts with n= 10-15 per group.

### Microbiota from aged mice is less resilient to an acute antibiotic challenge

Older adults are treated with antibiotics at higher rates (King et al., 2020). Moreover, while the microbiome appears largely resilient to acute antibiotic challenges in young populations, few studies have explored how composition and immunogenic capacity of the gut microbiota recovers from antibiotic usage during advanced age. Thus, we next compared how the gut microbiota from young and aged mice would respond to, and recover from, a broad-spectrum antibiotic (Abx) challenge. Young and aged mice were provided an Abx cocktail in drinking water for 7 days. We then analyzed immunogenic activity from feces immediately after Abx treatment (D0) (**Fig. 5**) as well as microbiome, host inflammatory status (**Fig. 4**), and immunogenicity (**Fig. 5**) at 7 days after the cessation of Abx treatment (D7).

**Figure 4.**
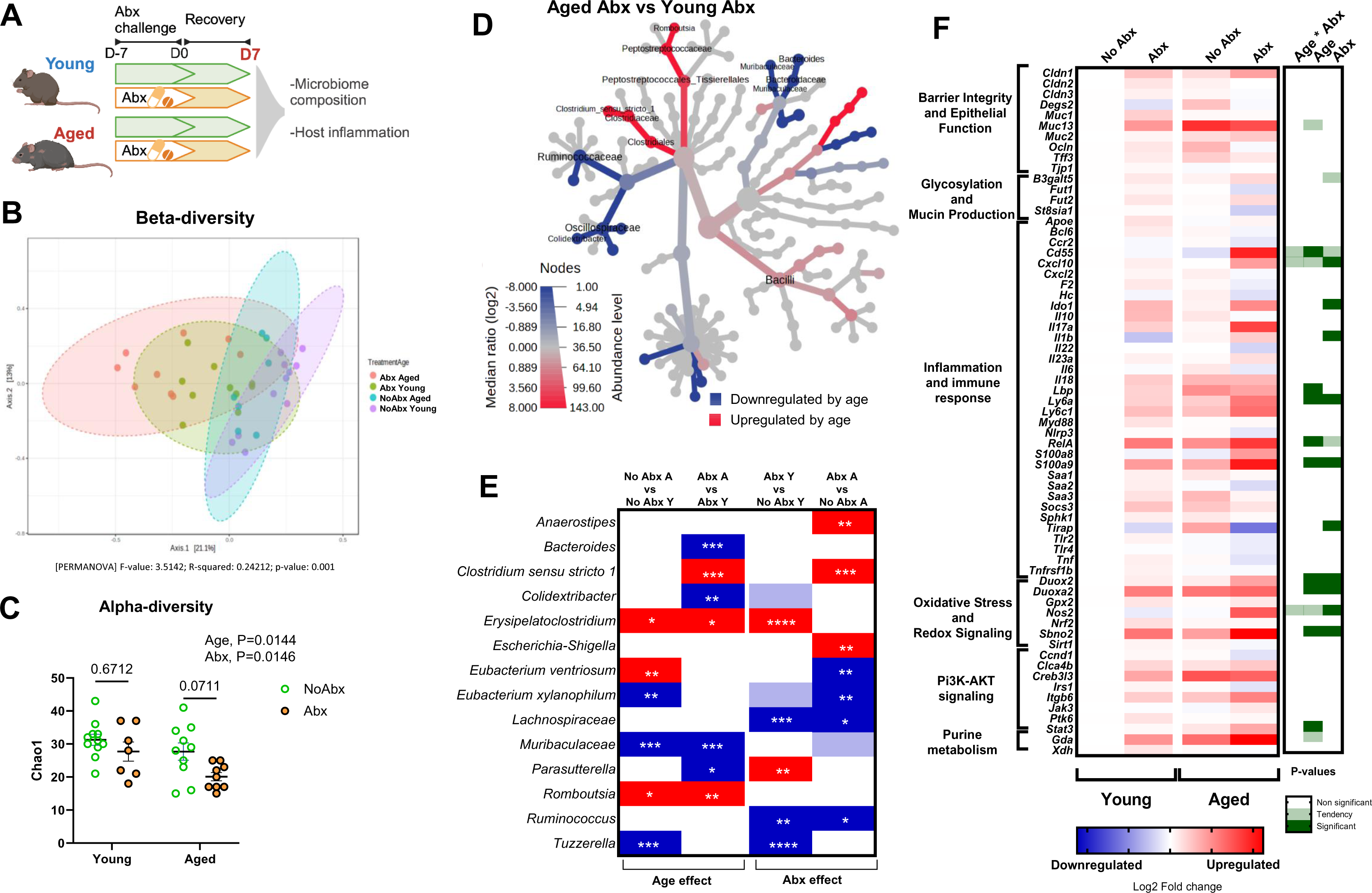
Age impacts gut microbiome and intestinal inflammation during recovery from treatment with antibiotics. **(A)** A schematic representation of antibiotic experiment. Mice were exposed to broad-spectrum antibiotics (Abx) in drinking water for seven (7) days and then were switched to regular drinking water for an additional seven days. Microbiome and gene expression analyses were carried out in proximal colon contents and tissue, respectively. All panels convey data from D7 (7 days after cessation of Abx treatment). **(B)** Microbiome β-diversity Bray-Curtis – Principal Coordinate Analysis (PcoA); dotted-line ellipses corresponding to clusters in each group. **(C)** Microbiome α-diversity – Chao 1 represents community richness in aged vs young mice exposed to antibiotics. **(D)** Heat tree representing taxa changes in aged vs young mice, both treated with antibiotics. **(E)** Bacterial genera significantly altered by age and/or antibiotics – Significant changes assessed by Kruskal-Wallis nonparametric test in taxa abundance. Dark red-significant increase, dark blue-significance decrease * *p*<0.05; ** *p*<0.01; *** *p*<0.001; **** *p*<0.0001. Light colors (blue or red) represent tendency (0.05<*p*≤0.1) and white indicates non-significant (*p*>0.1) changes in taxa abundance. Y = young, A = aged. Microbiome data correspond to n= 3 cohorts with n= 7-11 per group. **(F)** Heat map of distal colon gene expression analyzed via Fluidigm. Blue indicates downregulation and red indicates upregulation compared to young mice. Each square represents the mean gene expression per group (n = 8-12). The right panel visually conveys statistical significance: white implies no effect (*p*>0.10), light green represents a trend (0.05 <*p*≤ 0.10), and dark green indicates a significant change (*p*≤0.05). Gene expression data correspond to n= 3 cohorts with n= 12-15 per group.

**Figure 5.**
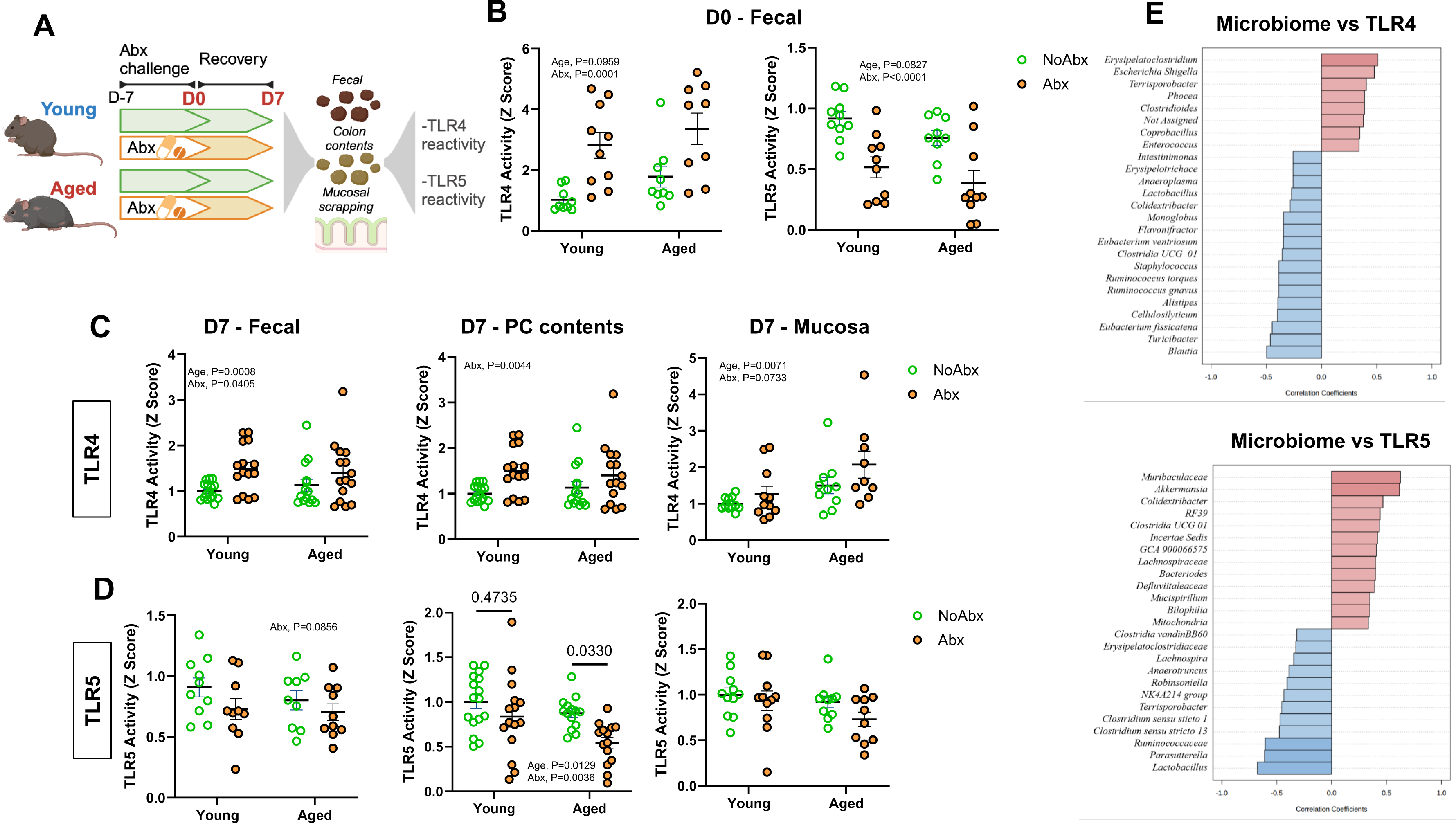
Age impacts microbiota-dependent immunogenic signatures after treatment with antibiotics. **(A)** A schematic representation of antibiotic experiment in immunogenicity. Mice were exposed to broad-spectrum antibiotics (Abx) in drinking water for seven (7) days and then were switched to regular drinking water for an additional seven days. **(B)** TLR4 and TLR5 activity in feces after Abx treatment (D0); **(C)** TLR 4 and **(D)** TLR5 activity in feces, proximal colon (PC) contents and mucosal scrapings after recovery period (D7), assessed using human embryonic kidney (HEK)-Blue-mTLR4 and HEK-Blue-mTLR5 cells. Data was normalized within each independent experiment and relativized to young control at AU =1. **(E)** Top 25 genera correlated with TLR4 and TLR5 activity (Spearman rank correlation). Data correspond to n= 3 cohorts with n= 9-15 per group.

As expected, substantial shifts in the microbiome composition of both young and aged mice were observed 7 days post Abx challenge (D7) (Bray-Curtis β-diversity PERMANOVA p<0.01; **Fig. 4B**). Yet, aged animals exhibited a unique response to Abx compared to young mice including a reduction in α-diversity (as measured by Chao1) in aged, but not young, mice (**Fig. 4C**). Abx treatment increased the abundance of *Erysipelatoclostridium* and *Parasuterella,* and decreased the abundance of members of *Lachnospiraceae*, *Ruminococcus*, and *Tuzzerella* in young mice. Conversely, Abx treatment increased *Anaerostipes*, *Clostridium*, and *Escherichia*, and decreased *Eubacterium xylanophilum, Lachnospiraceae* and *Ruminococcus* in aged mice (**Fig. 4E**, FDR *p*<0.05). We also observed unique colonic transcriptional signatures between aged and young mice in response to Abx, primarily in genes associated with inflammation and immune response, as well as oxidative stress and redox signaling (**Fig. 4F**). This included age and Abx-induced increases in key reactive oxygen (ROS) and reactive nitrogen species (RNS) producing enzymes *Duox2, Duoxa2,* and *Nos2* (**Fig. 4F**); dual-oxidase 2 (DUOX2) and its maturation factor (DUOXA2), as well as inducible nitric oxide synthase (iNOS) are critical for maintenance of redox tone in the distal bowel.

We next determined how age would impact microbiota immunogenicity in response to a broad-spectrum Abx challenge. We found that Abx exposure led to a robust age-independent increase in fecal TLR4 reactivity immediately post Abx challenge (D0) (**Fig. 5B**, *p*<0.01). This effect contrasted a robust decrease in the ability of feces of both age groups to activate TLR5 immediately after Abx (**Fig. 5B**, *p*<0.01). After a 7-day recovery period (D7), fecal (*p*<0.05), proximal colon (PC) contents (*p*<0.01), and mucosal scrapings (*p*=0.07) TLR4 reactivity remained elevated in both age groups (**Fig. 5C**). In contrast, while colonic contents and mucosal scrapings of young mice recovered TLR5 activity 7 days after Abx (*p*>0.05 vs young no Abx), aged mice exposed to antibiotics showed suppressed restoration of proximal colon content TLR5 immunogenicity (*p*<0.05 vs aged no Abx; **Fig. 5D)**.

To support the role of the microbiome as a driver of TLR-mediated immunogenicity, we analyzed Spearman correlations between microbial taxa and TLR4/TLR5 activity (**Fig. 5E**). We showed that TLR4 activity was associated with *Erysipelatoclostridium sp.* and *Escherichia sp.*, both of which were increased by age. *Colidextribacter sp.* and *Muribaculaceae sp.* were decreased by age, and positively correlated with TLR5, whereas *Clostridium sp.* were increased by age and negatively associated with TLR5.

## DISCUSSION

The gut microbiota changes throughout the host’s lifespan and is associated with various aspects of aging physiology, including chronic low-grade inflammation. However, the functional relationships between the gut microbiota and inflammatory processes during advanced age are still not fully understood. Here, we unraveled a previously unexplored role of gut microbiota immunogenic signatures within the distal bowel that are tightly linked to host inflammatory tone with advanced aged.

This study first corroborated the well-established effect of aging on microbiome composition (DeJong et al., 2020). The gut microbiota of aged mice was markedly reduced in microbial diversity and shifts in microbial taxa that strongly corresponded to host inflammation, indicating signs of dysbiosis associated with aging (Petersen & Round, 2014). Our findings are in agreement with previous studies (Hodgkinson et al., 2023; Lee et al., 2020; Vital et al., 2014), which identified an age-induced decline in obligate anaerobes with known butyrate producing capacity, such as members of *Lachnospiraceae* and *Ruminococcaceae* families.

Previous studies have also identified the gut microbiota to be associated with an increased inflammatory status during aging. This has been evidenced by increased systemic inflammatory markers (Biagi et al., 2010), greater intestinal permeability, and decreased macrophage function (Thevaranjan et al., 2017), which are linked to concomitant shifts in the microbiome. We corroborated and expanded on these findings by showing that an aging microbiota is tightly associated with signs of barrier disruption and innate inflammatory infiltration, including robust elevations in LBP and calprotectin, respectively. The cytoplasmic calprotectin is found in innate immune cells, including neutrophils and monocytes and is a gold standard biomarker used for assessing ongoing inflammatory processes in the context of inflammatory bowel diseases (Fernandes et al.; Ling Lundström et al., 2023). While largely understudied in the context of aging, our findings in mice corroborates data from aged humans showing that fecal calprotectin progressively increases with age, especially within individuals over 60 years old of age (Joshi et al., 2010).

We showed for the first time that advanced age results in increased capacity for both colonic contents and mucosal scrapings to activate a critical inflammatory receptor, TLR4. Our findings regarding age-associated increases in microbiota-driven TLR4 immunogenicity corroborate other studies that have indicated alterations in host TLR4 signaling pathways that occur with age. For example, Bailey et al. (2019) has shown that aging dysregulates the innate immune response to TLR4 ligands in humans and leads to increased susceptibility to sepsis. Our work also corroborates recent work which showed that the gut microbiota LPS fraction of aged mice directly increases the inflammatory response in macrophages in a TLR4-dependent fashion (Kim et al., 2016). Nevertheless, even though TLR4 primarily recognizes bacterial LPS, certain endogenous molecules, including specific saturated fatty acids such as lauric acid (Hwang et al., 2016), high-mobility group box-1 (HMGB-1), heat shock proteins (HSPs) and others (Karuppagounder et al., 2016) may also activate TLR4 (Yousefi-Ahmadipour et al., 2022). Thus, our results suggest the possibility of a combined effect for microbiota LPS and other immunogenic molecules in the gut environment that enhances TLR4 activity, but this remains to be tested.

We also provided evidence that transfer of microbiota from aged mice could elicit age-associated inflammatory phenotypes in young germ-free recipients, suggesting that microbiota-driven TLR4 overactivation may have a direct role in shaping the pro-inflammatory environment through factors such as IL6 and TNF. This corroborates previous evidence showing that a microbiota transfer from aged mice to young germ-free mice contributes to heightened gut and systemic inflammation. For example, the microbiota from aged mice has been reported to have obesogenic properties (Binyamin et al., 2020), exacerbate inflammatory markers including IL-6, IL-1β, TNF-α (Crossland et al., 2023), and increase translocation of inflammatory bacterial products into the circulation of germ-free recipients (Fransen et al., 2017).

Older adults are exposed to antibiotics at higher rates as compared to young individuals. This is critical because older adults have higher rates of infection (Ciarambino et al., 2023) and antibiotic-induced perturbations to the gut microbiota have been directly linked with reduced vaccine efficacy in older adults who already suffer from reduced responses to vaccine (Ciarambino et al., 2023; Hagan et al., 2019). Our findings revealed notable variations in recovery from antibiotics in aged vs. young mice. This included taxa such as *Clostridium sp., Erysipelatoclostridium sp., Eubacterium ventriosum* and *Escherichia sp.*, that responded differentially to antibiotic treatment in aged mice.

Further, we showed that age-associated attenuation in gut microbiota recovery from an antibiotic challenge paralleled changes in colonic inflammatory status, inducing genes encoding dual-oxidase 2 (DUOX2) and inducible nitric oxide synthase (iNOS; also known as NOS2), two critical proteins that control redox tone in the distal bowel (Aviello & Knaus, 2018; Mu et al., 2019). Altered redox balance in the gut is critical for shaping the gut microbiota composition. Higher concentrations of host-derived nitrate and ROS (driven by genes such as *Nos2* and *Duox2*) lead to the expansion of bacterial taxa that can selectively survive in a high ROS/RNS niche like *Escherichia*, which is resistant to ROS and can use nitrate as an electron acceptor (Byndloss et al., 2017; Winter et al., 2013). The observed concurrent elevations of *Escherichia sp.* and colonic *Nos2* and *Duox2* in aged mice exposed to antibiotics suggests that elevated ROS activity in the aging gut is a potential mediator of microbiota perturbations.

Aged mice showed a more pronounced reduction in TLR5 activity during recovery after treatment with antibiotics. In contrast to TLR4, TLR5 activation by the gut microbiota is recognized for its capacity to suppress inflammation, promote tissue regeneration in models of disease, and strengthen gut barrier function (Lim et al., 2024). Moreover, previous studies have indicated that antibiotics decrease TLR5-mediated flagellin sensing, leading to a reduction in vaccine efficacy (Oh et al., 2014). Our findings extend this observation by demonstrating that antibiotics robustly impact TLR5 immunogenicity in aged mice versus their young counterparts and underscore the microbiota as a potential promotor of immunosenescence in the distal gut.

Collectively, our data point to unique immunogenic signatures of the aged gut microbiota, underscored by enhanced TLR4 immunogenicity that is directly linked to inflammation of the colon. We also provide evidence that a microbiota of aged mice is less resilient to environmental challenges, as highlighted by a reduced ability to recover microbiota immunogenic properties following exposure to broad-spectrum antibiotics. Future studies will define the immunogenic capacity of the microbiota as a regulator of age-associated inflammation and chronic disease.

## METHODS

### Animals

Young (8-10 months) and aged (18-20 months) conventionally raised (CONV-R) C57BL/6 mice were provided through National Institutes of Aging (NIA) SPF aging colony. Mice were shipped to and maintained at University of Illinois Urbana-Champaign under institutionally approved protocol (IACUC #22203). After a two-week acclimation period, young and aged mice were assessed across independent cohorts, balancing male and female. The number of cohorts and n per group are described in the Figure legends. Mice were housed with 3-4 mice per cage with *ad libitum* access to water and regular chow diet throughout the experiments. A light:dark 12 h cycle was used, with lights on at 6 AM and off at 6 PM.

#### Germ-free (GF) mice

Three (3) to four (4) month-old germ-free mice were obtained from The Ohio State University Gnotobiotic and Germ-Free Mouse Resource. To prevent microbial contamination, germ-free animals were kept in sentry sealed positive pressure cages (Allentown, Allentown, NJ) for a 1-week acclimation period and throughout the course of the study. After the acclimation period, the protocol from the National Gnotobiotic Rodent Resource Center (NGRRC) was utilized to confirm germ-free sterility in such mice at the Research Institute at National wide Children’s Hospital (Columbus, OH). Microbiota transplantation (MT) was prepared utilizing a weight standardized mixture of intestinal contents (cecal contents, colonic contents and feces) from young and aged CONV-R mice. Frozen fecal matter was added to anaerobic PBS (1:4 weight to volume) and then homogenized by pipetting without introducing air bubbles. The sample was then centrifuged at 500 × *g* for 1 min to obtain the supernatant. MT to germ-free mice from young (n = 10) and aged (n = 15) CONV-R mice was performed by administering an aliquot of supernatant (100 µL) by oral gavage. Mice were euthanized by CO_2_ asphyxiation 4 weeks after MT.

#### Antibiotics treatment

The study with antibiotic (**Abx**) treatment was performed by providing a broad-spectrum antibiotic cocktail (neomycin 40 mg/kg/day, ampicillin 33 mg/kg/day, metronidazole 21.5 mg/kg/day, vancomycin 4.5 mg/kg/day; Sigma-Aldrich) in drinking water for 7 days to the assigned mice for each group. Control mice (**No Abx**) received regular water throughout the experiment. Abx did not affect water consumption.

Feces were collected from all mice immediately after completion of Abx treatment (D0), and 7 days after cessation of Abx treatment (D7). After the sacrifice by CO_2_ asphyxiation at D7, colon length, colon weight, spleen weight and cecum weight were measured. Serum and large intestine were collected and stored at −80°C for further analysis.

### 16s rRNA gene sequencing

Colon contents (∼10 mg) were incubated for 45 min at 37 °C in lysozyme buffer (22 mg/ml lysozyme, 20 mM TrisHCl, 2 mM EDTA, 1.2% Triton-x, pH 8.0) before homogenization for 150 s with 0.1 mm zirconia beads. Then, samples were incubated at 95 °C for 5 min with InhibitEX Buffer, followed by incubation at 70 °C for 10 min with Proteinase K and lysis Buffer AL. Next, QIAamp Fast DNA Stool Mini Kit (Cat # 51604, Qiagen, Hilden Germany) was utilized to extract DNA from the homogenized content. All the conditions followed manufacturer’s instructions, with slight modifications as previously described by Allen et al. (2022). dsDNA Broad Range Assay Kit was used to quantify DNA with Qubit 2.0 Fluorometer (Life Technologies, Carlsbad, CA). Illumina MiSeq was used to obtain amplified PCR libraries sequencing done from both ends of 250 nt region of V4 16S rRNA hypervariable region. Amplicon processing and downstream taxonomic assignment using the ribosomal RNA database SILVA v138 was performed using DADA2 and QIIME 2.0 platform. EMPeror was used to visualize the microbial diversity (β-diversity, Bray-Curtis) in 3-dimensional PCoA plots.

### TLR4 and TLR5 reporter assays

Assessment of TLR4 and TLR5 activity was implemented using human embryonic kidney (HEK)-Blue-mTLR4 and HEK-Blue-mTLR5 cells respectively (Invivogen) using a protocol as previously described (Allen et al., 2022; Tran et al., 2019). Briefly, feces, intestinal contents or mucosal scrapings were resuspended in PBS w/v (100 mg/mL) and homogenized for 5 min using vortex. Each sample was then centrifuged at 8000 × *g* for 2 min, serially diluted, and supernatant applied directly to either HEK-Blue-mTLR4 or HEK-Blue-mTLR5 cells (2 x 10^5^cells/well). After 18h of stimulation, cell culture supernatants were applied to QUANTI-Blue medium (Invivogen) and measured for alkaline phosphatase activity at 620 nm after 90 mins. Relative fluorescence intensity was used as an estimation for TLR4 or TLR5 activity, considering young mice no Abx as control (Relative AU =1).

### Gene Expression by Fluidigm

Colon tissue RNA was isolated using the PureLink RNA Mini Kit (Cat # 12183018A, Thermo Fisher Scientific) and cDNA synthesized with a High-Capacity cDNA Reverse Transcription Kit (Cat # 4368814, Thermo Fisher Scientific). Primers (Fwd and Rv, Integrated DNA Technologies, Coralville, IA) used are listed in Table S1. Real-Time PCR Analysis was performed by the University of Illinois at Urbana-Champaign Functional Genomics Unit of the W.M. Keck Center by Fluidigm analysis (96 × 96 chip) technology. The software Fluidigm Real-Time PCR Analysis 3.0.2 (Fluidigm, San Francisco, CA, USA) was used to acquire the data. Relative expression was determined using the delta-delta cycle threshold method (dd*C*_t_) using Young No Abx mice as control. Eukaryotic elongation factor 2 (*Eef2*) was used as housekeeping gene. Values were log2 transformed prior to statistical analysis.

### Serum LBP and Colon Content Calprotectin

Commercially available ELISA kits (Hycult Biotec, Uden, Netherlands) were utilized according to manufacturers’ instructions to assess LPS-binding protein (LBP) and calprotectin concentrations in serum and colon contents, respectively.

### Statistical analyses

All data was expressed as mean ± SEM. Downstream Microbiome statistics were performed using the Microbiome Analyst software (http://www.microbiomeanalyst.ca) (Lu et al., 2023). Data was rarefied to the minimum library size and transformed using the center log ratio (CLR) method to mitigate compositional biases. The Bray-Curtis dissimilarity index (β-diversity) was implemented to assess community compositional differences between samples and assessed via Permutational multivariate analysis of variance (PERMANOVA). Principal Coordinate Analysis (PCoA) was utilized for visualization. Chao1-index was utilized to assess ⍺-diversity. Effects of age and antibiotics on the gut microbiota was assessed by multiple linear regression with covariate adjustment using a linear model and adjusted p<0.05. Taxonomic data was visualized with heat trees (Foster et al., 2017). Spearman rank correlation was used to analyze associations between microbiome taxonomic/immunogenicity and host inflammatory outcomes.

To account for differences in experimental cohorts, data were Z-scored relative to the control group within each independent cohort. Student T-tests or Non-parametric Mann-Whitney U tests were used to compare differences in immunogenicity and inflammatory outcomes in aged vs. young mice. Two-way analysis of variance (ANOVA) was implemented to examine age and antibiotic effects on host inflammatory signatures. The statistical analyses were performed using the statistical package GraphPad Prism 8.0.1 (GraphPad Software, Inc., San Diego, CA, USA) with a priori ⍺ set at 0.05. The Benjamini-Hochberg method for controlling false discovery rate (FDR) was implemented to limit chance of type-1 errors in multiple testing analyses.

## Author contributions

TWB, JMA conceived and designed the study. MTB, JAW, TWB, JMA provided funding and resources. MECS, AS, AFM, DK, PCB, MCK, CHL, DHS, MTB, and JMA conducted the experiments and acquired the data. MECS, AS, PCB, MCK, MTB, and JMA analyzed and interpreted the data. MECS, AS, and JMA drafted the manuscript. MECS, AS, AFM, DK, PCB, MCK, DAW, DHS, JAW, MTB, TWB, and JMA critically revised the manuscript. MECS, AS, AFM, DK, PCB, MCK, CHL, DAW, DHS, JAW, MTB, TWB, and JMA approved the final version to be submitted for publication.

## Data Availability Statement

All data that support the findings of this study are available from the authors upon request. Microbiome data will be made public under NCBI upon acceptance of publication.

## Acknowledgments

This work was supported by the National Institutes of Health (NIH). TWB, JAW, MTB and JMA were supported by R56AG068747 (NIH/NIA). JMA was supported by R01DK131133 (NIH/NIDDK).

## References

Allen, J. M., Mackos, A. R., Jaggers, R. M., Brewster, P. C., Webb, M., Lin, C.-H., Ladaika, C., Davies, R., White, P., & Loman, B. R. (2022). Psychological stress disrupts intestinal epithelial cell function and mucosal integrity through microbe and host-directed processes. Gut Microbes, 14(1), 2035661.

Alsegiani, A. S., & Shah, Z. A. (2022). The influence of gut microbiota alteration on age-related neuroinflammation and cognitive decline. Neural Regen Res, 17(11), 2407–2412. 10.4103/1673-5374.335837

Aviello, G., & Knaus, U. G. (2018). NADPH oxidases and ROS signaling in the gastrointestinal tract. Mucosal Immunology, 11(4), 1011–1023. 10.1038/s41385-018-0021-8

Bailey, K. L., Smith, L. M., Heires, A. J., Katafiasz, D. M., Romberger, D. J., & LeVan, T. D. (2019). Aging leads to dysfunctional innate immune responses to TLR2 and TLR4 agonists. Aging Clin Exp Res, 31(9), 1185–1193. 10.1007/s40520-018-1064-0

Biagi, E., Nylund, L., Candela, M., Ostan, R., Bucci, L., Pini, E., Nikkïla, J., Monti, D., Satokari, R., Franceschi, C., Brigidi, P., & De Vos, W. (2010). Through ageing, and beyond: gut microbiota and inflammatory status in seniors and centenarians. PLoS One, 5(5), e10667. 10.1371/journal.pone.0010667

Binyamin, D., Werbner, N., Nuriel-Ohayon, M., Uzan, A., Mor, H., Abbas, A., Ziv, O., Teperino, R., Gutman, R., & Koren, O. (2020). The aging mouse microbiome has obesogenic characteristics. Genome Medicine, 12(1), 87. 10.1186/s13073-020-00784-9

Byndloss, M. X., Olsan, E. E., Rivera-Chávez, F., Tiffany, C. R., Cevallos, S. A., Lokken, K. L., Torres, T. P., Byndloss, A. J., Faber, F., & Gao, Y. (2017). Microbiota-activated PPAR-γ signaling inhibits dysbiotic Enterobacteriaceae expansion. Science, 357(6351), 570–575.

Ciarambino, T., Crispino, P., Buono, P., Giordano, V., Trama, U., Iodice, V., Leoncini, L., & Giordano, M. (2023). Efficacy and Safety of Vaccinations in Geriatric Patients: A Literature Review. Vaccines, 11(9), 1412. https://www.mdpi.com/2076-393X/11/9/1412

Conway, J., & A Duggal, N. (2021). Ageing of the gut microbiome: Potential influences on immune senescence and inflammageing. Ageing Research Reviews, 68, 101323. 10.1016/j.arr.2021.101323

Crossland, N. A., Beck, S., Tan, W. Y., Lo, M., Mason, J. B., Zhang, C., Guo, W., & Crott, J. W. (2023). Fecal microbiota transplanted from old mice promotes more colonic inflammation, proliferation, and tumor formation in azoxymethane-treated A/J mice than microbiota originating from young mice. Gut Microbes, 15(2), 2288187. 10.1080/19490976.2023.2288187

DeJong, E. N., Surette, M. G., & Bowdish, D. M. (2020). The gut microbiota and unhealthy aging: disentangling cause from consequence. Cell host & microbe, 28(2), 180–189.

Fernandes, S. R., Bernardo, S., Saraiva, S., Gonçalves, A. R., Moura Santos, P., Valente, A., Correia, L. A., Cortez-Pinto, H., & Magro, F. Tight control using fecal calprotectin and early disease intervention increase the rates of transmural remission in Crohn’s disease. United European Gastroenterology Journal, n/a(n/a). 10.1002/ueg2.12497

Foster, Z. S. L., Sharpton, T. J., & Grünwald, N. J. (2017). Metacoder: An R package for visualization and manipulation of community taxonomic diversity data. PLOS Computational Biology, 13(2), e1005404. 10.1371/journal.pcbi.1005404

Fransen, F., van Beek, A. A., Borghuis, T., Aidy, S. E., Hugenholtz, F., van der Gaast-de Jongh, C., Savelkoul, H. F. J., De Jonge, M. I., Boekschoten, M. V., Smidt, H., Faas, M. M., & de Vos, P. (2017). Aged Gut Microbiota Contributes to Systemical Inflammaging after Transfer to Germ-Free Mice. Front Immunol, 8, 1385. 10.3389/fimmu.2017.01385

Ghosh, T. S., Shanahan, F., & O’Toole, P. W. (2022). The gut microbiome as a modulator of healthy ageing. Nature Reviews Gastroenterology & Hepatology, 19(9), 565–584. 10.1038/s41575-022-00605-x

Graham, D. B., & Xavier, R. J. (2023). Conditioning of the immune system by the microbiome. Trends Immunol, 44(7), 499–511. 10.1016/j.it.2023.05.002

Hagan, T., Cortese, M., Rouphael, N., Boudreau, C., Linde, C., Maddur, M. S., Das, J., Wang, H., Guthmiller, J., Zheng, N.-Y., Huang, M., Uphadhyay, A. A., Gardinassi, L., Petitdemange, C., McCullough, M. P., Johnson, S. J., Gill, K., Cervasi, B., Zou, J.,…Pulendran, B. (2019). Antibiotics-Driven Gut Microbiome Perturbation Alters Immunity to Vaccines in Humans. Cell, 178(6), 1313–1328.e1313. 10.1016/j.cell.2019.08.010

Hodgkinson, K., El Abbar, F., Dobranowski, P., Manoogian, J., Butcher, J., Figeys, D., Mack, D., & Stintzi, A. (2023). Butyrate’s role in human health and the current progress towards its clinical application to treat gastrointestinal disease. Clinical Nutrition, 42(2), 61–75. 10.1016/j.clnu.2022.10.024

Hwang, D. H., Kim, J.-A., & Lee, J. Y. (2016). Mechanisms for the activation of Toll-like receptor 2/4 by saturated fatty acids and inhibition by docosahexaenoic acid. European Journal of Pharmacology, 785, 24–35. 10.1016/j.ejphar.2016.04.024

Joshi, S., Lewis, S. J., Creanor, S., & Ayling, R. M. (2010). Age-related faecal calprotectin, lactoferrin and tumour M2-PK concentrations in healthy volunteers. Ann Clin Biochem, 47(Pt 3), 259–263. 10.1258/acb.2009.009061

Kabbani, S., Palms, D., Bartoces, M., Stone, N., & Hicks, L. A. (2018). Outpatient Antibiotic Prescribing for Older Adults in the United States: 2011 to 2014. J Am Geriatr Soc, 66(10), 1998–2002. 10.1111/jgs.15518

Karuppagounder, V., Giridharan, V. V., Arumugam, S., Sreedhar, R., Palaniyandi, S. S., Krishnamurthy, P., Quevedo, J., Watanabe, K., Konishi, T., & Thandavarayan, R. A. (2016). Modulation of Macrophage Polarization and HMGB1-TLR2/TLR4 Cascade Plays a Crucial Role for Cardiac Remodeling in Senescence-Accelerated Prone Mice. PLoS One, 11(4), e0152922. 10.1371/journal.pone.0152922

Kim, K.-A., Jeong, J.-J., Yoo, S.-Y., & Kim, D.-H. (2016). Gut microbiota lipopolysaccharide accelerates inflamm-aging in mice. BMC microbiology, 16, 1–9.

King, L. M., Bartoces, M., Fleming-Dutra, K. E., Roberts, R. M., & Hicks, L. A. (2020). Changes in US Outpatient Antibiotic Prescriptions From 2011-2016. Clin Infect Dis, 70(3), 370–377. 10.1093/cid/ciz225

Laubitz, D., Typpo, K., Midura-Kiela, M., Brown, C., Barberan, A., Ghishan, F. K., & Kiela, P. R. (2021). Dynamics of Gut Microbiota Recovery after Antibiotic Exposure in Young and Old Mice (A Pilot Study). Microorganisms, 9(3). 10.3390/microorganisms9030647

Lee, J., Venna, V. R., Durgan, D. J., Shi, H., Hudobenko, J., Putluri, N., Petrosino, J., McCullough, L. D., & Bryan, R. M. (2020). Young versus aged microbiota transplants to germ-free mice: increased short-chain fatty acids and improved cognitive performance. Gut Microbes, 12(1), 1814107. 10.1080/19490976.2020.1814107

Lim, J. S., Jeon, E. J., Go, H. S., Kim, H.-J., Kim, K. Y., Nguyen, T. Q. T., Lee, D. Y., Kim, K. S., Pietrocola, F., & Hong, S. H. (2024). Mucosal TLR5 activation controls healthspan and longevity. Nature Communications, 15(1), 46.

Ling Lundström, M., Peterson, C., Lampinen, M., Hedin, C. R. H., Keita Å, V., Kruse, R., Magnusson, M. K., Lindqvist, C. M., Repsilber, D., D’Amato, M., Hjortswang, H., Strid, H., Rönnblom, A., Söderholm, J. D., Öhman, L., Venge, P., Halfvarson, J., & Carlson, M. (2023). Fecal Biomarkers of Neutrophil and Eosinophil Origin Reflect the Response to Biological Therapy and Corticosteroids in Patients With Inflammatory Bowel Disease. Clin Transl Gastroenterol, 14(8), e00605. 10.14309/ctg.0000000000000605

Lu, Y., Zhou, G., Ewald, J., Pang, Z., Shiri, T., & Xia, J. (2023). MicrobiomeAnalyst 2.0: comprehensive statistical, functional and integrative analysis of microbiome data. Nucleic Acids Res, 51(W1), W310–w318. 10.1093/nar/gkad407

Malik, J. A., Zafar, M. A., Lamba, T., Nanda, S., Khan, M. A., & Agrewala, J. N. (2023). The impact of aging-induced gut microbiome dysbiosis on dendritic cells and lung diseases. Gut Microbes, 15(2), 2290643. 10.1080/19490976.2023.2290643

Mu, K., Yu, S., & Kitts, D. D. (2019). The Role of Nitric Oxide in Regulating Intestinal Redox Status and Intestinal Epithelial Cell Functionality. Int J Mol Sci, 20(7). 10.3390/ijms20071755

Oh, J. Z., Ravindran, R., Chassaing, B., Carvalho, F. A., Maddur, M. S., Bower, M., Hakimpour, P., Gill, K. P., Nakaya, H. I., Yarovinsky, F., Sartor, R. B., Gewirtz, A. T., & Pulendran, B. (2014). TLR5-mediated sensing of gut microbiota is necessary for antibody responses to seasonal influenza vaccination. Immunity, 41(3), 478–492. 10.1016/j.immuni.2014.08.009

Park, S. Y. (2020). Age-related fecal calprotectin concentrations in healthy adults. Korean Journal of Clinical Laboratory Science, 52(3), 181–187.

Petersen, C., & Round, J. L. (2014). Defining dysbiosis and its influence on host immunity and disease. Cellular microbiology, 16(7), 1024–1033.

Sakurai, T., & Saruta, M. (2022). Positioning and Usefulness of Biomarkers in Inflammatory Bowel Disease. Digestion, 104(1), 30–41. 10.1159/000527846

Thevaranjan, N., Puchta, A., Schulz, C., Naidoo, A., Szamosi, J., Verschoor, C. P., Loukov, D., Schenck, L. P., Jury, J., & Foley, K. P. (2017). Age-associated microbial dysbiosis promotes intestinal permeability, systemic inflammation, and macrophage dysfunction. Cell host & microbe, 21(4), 455–466. e454.

Tran, H. Q., Ley, R. E., Gewirtz, A. T., & Chassaing, B. (2019). Flagellin-elicited adaptive immunity suppresses flagellated microbiota and vaccinates against chronic inflammatory diseases. Nature communications, 10(1), 5650.

Vital, M., Howe, A. C., & Tiedje, J. M. (2014). Revealing the Bacterial Butyrate Synthesis Pathways by Analyzing (Meta)genomic Data. mBio, 5(2), 10.1128/mbio.00889-00814. 10.1128/mbio.00889-14

Wilmanski, T., Diener, C., Rappaport, N., Patwardhan, S., Wiedrick, J., Lapidus, J., Earls, J. C., Zimmer, A., Glusman, G., Robinson, M., Yurkovich, J. T., Kado, D. M., Cauley, J. A., Zmuda, J., Lane, N. E., Magis, A. T., Lovejoy, J. C., Hood, L., Gibbons, S. M.,…Price, N. D. (2021). Gut microbiome pattern reflects healthy ageing and predicts survival in humans. Nat Metab, 3(2), 274–286. 10.1038/s42255-021-00348-0

Winter, S. E., Winter, M. G., Xavier, M. N., Thiennimitr, P., Poon, V., Keestra, A. M., Laughlin, R. C., Gomez, G., Wu, J., & Lawhon, S. D. (2013). Host-derived nitrate boosts growth of E. coli in the inflamed gut. Science, 339(6120), 708–711.

Yousefi-Ahmadipour, A., Sartipi, M., Khodadadi, H., Shariati-Kohbanani, M., & Arababadi, M. K. (2022). Toll-like receptor 4 and the inflammation during aging. JOURNAL OF GERONTOLOGY AND GERIATRICS, 70, 144–151.

Zhu, X., Li, B., Lou, P., Dai, T., Chen, Y., Zhuge, A., Yuan, Y., & Li, L. (2021). The Relationship Between the Gut Microbiome and Neurodegenerative Diseases. Neurosci Bull, 37(10), 1510–1522. 10.1007/s12264-021-00730-8

